# USP20 deubiquitinates and stabilizes the ER-phagy receptor FAM134B to drive ER-phagy

**DOI:** 10.1101/2023.07.27.550606

**Authors:** Zhang Man, Zhangshun Wang, Qing Zhao, Qian Yang, Cuiwei Yang, Yanfen Liu

**Affiliations:** School of Life Science and Technology, ShanghaiTech University, Shanghai 201210, China

**Keywords:** deubiquitination, ER-phagy, FAM134B, LC3, USP20, VAPs

## Abstract

The endoplasmic reticulum (ER) serves as a hub for various essential cellular processes, and maintaining ER homeostasis is essential for cell function. ER-phagy is a selective process that removes impaired ER subdomains through autophagosomes and lysosomal degradation. While the involvement of ubiquitination in autophagy regulation is well-established, its role in ER-phagy remains unclear. In our study, we screened deubiquitinating enzymes involved in ER-phagy and identified USP20 (ubiquitin specific peptidase 20) as a key regulator of ER-phagy under stress conditions. USP20 specifically cleaves K63– and K48-linked ubiquitin chains on the ER-phagy receptor FAM134B/RETREG1 (reticulophagy regulator 1), thereby stabilizing the substrate and promoting ER-phagy. Remarkably, despite lacking a transmembrane domain, USP20 is recruited to the ER through its interaction with VAPs (vesicle-associated membrane proteins). VAPs facilitate the recruitment of early autophagy proteins, including WIPI2, to specific ER subdomains, where USP20 and FAM134B are enriched. This recruitment of WIPI2 and other proteins plays a crucial role in facilitating FAM134B-mediated ER-phagy in response to cellular stress. Our findings highlight the critical role of USP20 in maintaining ER homeostasis by deubiquitinating and stabilizing FAM134B at distinct ER subdomains, where USP20 further recruits VAPs and promotes efficient ER-phagy.

## Introduction

Autophagy is a conservative eukaryotic cellular process that delivers protein aggregates, damaged organelles and invaded pathogens to lysosome for degradation [1–3]. It is essential for survival, differentiation, development and homeostasis, defects of which are linked to various diseases such as neurodegenerative diseases, inflammatory disorders and cancers [4, 5]. Autophagy is initiated by the formation of a specific double-membrane organelle called the autophagosome, which engulfs cargo to lysosome for destruction. It is a multistep process involving several key ATG proteins and signaling complexes. Among them, the ULK1/2 complex, consisting of ULK1/2 kinase, ATG13, RB1CC1/FIP200, and ATG101, is essential for autophagy initiation [6–8].

The endoplasmic reticulum (ER) is a membrane-bound organelle consisting of the nuclear envelope and the peripheral ER that contains sheets and tubules. The integrity of ER is essential for protein and lipid synthesis, protein quality control, calcium homeostasis and inter-organelle communication [9, 10]. In multicellular organisms, the ER serves as a platform where ATG proteins are recruited for the formation of autophagosomes [11]. The ER-resident tail-anchored VAP proteins, VAPA and VAPB, use their MSP domain to interact with the FFAT (two phenylalanines [FF] in an acidic tract) motif present in both RB1CC1/FIP200 and ULK1. These interactions are crucial for the recruitment of ULK1 and the stabilization of the ULK1 complex at the sites where autophagosomes are formed on the ER [12]. VAPA/B also interact with the WD40 repeat-containing PI(3)P-binding protein WIPI2 to tether the ER/isolation membrane for isolation membrane expansion [12]. The ER is not only a key site for the initiation of macroautophagy but also a substrate of autophagy [11, 13]. To maintain ER homeostasis, certain ER fractions can be selectively degraded through a pathway known as ER-phagy [13–15]. Mutations in genes encoding ER-phagy components have been linked to specific neurodegenerative diseases [16, 17].

Macro-ER-phagy relies on the activation of the ER-phagy receptors, which bridge damaged ER fractions and core autophagy components. Multiple ER-phagy receptors, including FAM134B, RTN3L, ATL3, SEC62, CCPG1, TEX264, and CALCOCO1, have been reported in mammals [18–25]. These receptors play distinct roles in recognizing and targeting specific ER subdomains for degradation. Most ER-phagy receptors contain transmembrane domains that locate them on the ER, while only CALCOCO1 locates in the cytosol but can interact with ER-resident proteins VAPs to mediate ER fraction degradation [25]. These receptors contain at least one LC3 (or GABARAP)-interacting region (LIR or GIR) that can bind to Atg8 family proteins. FAM134B is the first identified receptor responsible for the degradation of sheet ER. It contains reticulon homology domains (RHDs) that comprise two short hairpin-like transmembrane domains, enabling the generation of membrane curvature [18, 26]. It has an LIR motif in the C-terminus to interact with LC3. Recent research has revealed that FAM134B undergoes oligomerization prior to binding with LC3 [27]. Additionally, the oligomerization process of FAM134B is facilitated by phosphorylation and acetylation of RHDs [28]. However, detail molecular mechanisms governing ER-phagy under stress remain insufficiently understood.

Ubiquitination is a critical post-translational modification involved in multiple cellular pathways [29, 30]. The ubiquitination process is ATP-dependent and occurs through a cascade of enzymatic steps involving an E1 to activate ubiquitin, an E2 to conjugate ubiquitin, and an E3 to facilitate the attachment of ubiquitin to the target protein [29]. Different types of polyubiquitin chains mediate distinct signaling pathways, ultimately determining the fate of the substrate protein [29, 30]. Conversely, deubiquitinating enzymes (DUBs) catalyze the removal of ubiquitin from target proteins, allowing for precise temporal and spatial control of protein ubiquitination [31]. E3 ligases and DUBs work in concert to regulate the ubiquitin chain assembly on specific substrates, thereby influencing various biological processes in organisms. Recent studies have shown its close association with autophagy, influencing the regulation of various autophagy-related proteins and impacting the autophagy flux [32–40]. Whether ubiquitination regulates ER-phagy has not been cleared elucidated.

In this study, we screened a DUB library and identified USP20 as a key factor regulating ER-phagy under stress. USP20 specifically cleaves K63 and K48-formed ubiquitin chain conjugated on the ER-phagy receptor FAM134B. The deubiquitination of FAM134B results in protein stabilization, which subsequently drives ER-phagy to resolve ER stress. Despite lacking a transmembrane domain, USP20 is consistently recruited to the ER through its interaction with ER-resident proteins known as VAPs. The interaction of USP20 with VAPs facilitates VAPs localization to specific ER subdomains under stress, where USP20 forms puncta with the ER-phagy receptor FAM134B. Concurrently, VAPs recruit autophagy initiation proteins including WIPI2 to these subdomains, further enhancing ER-phagy. These findings underscore the pivotal role of USP20 in promoting ER-phagy through the deubiquitination of FAM134B at specific ER subdomains, as well as in facilitating the assembly of autophagy initiation proteins, thereby contributing to the maintenance of ER homeostasis.

## Results

### USP20 promotes autophagy at multiple steps

To investigate the role of ubiquitin-dependent regulation in autophagy, we conducted an unbiased screen by individually overexpressing each of the DUBs in cells and assessing the changes in LC3B. Among the candidate DUBs, we were particularly interested in *USP20*, as its overexpression resulted in an elevated ratio of LC3B-II to LC3B-I compared to the control under starvation conditions using EBSS medium (**Fig. S1A-B**). The mRFP-EGFP-LC3B assay serves as an autophagic indicator, enabling the characterization of different stages of the autophagy pathway. It allows the visualization of autophagosomes (RFP^+^GFP^+^, appearing as yellow puncta) and autolysosomes (RFP^+^GFP^-^, appearing as red puncta) due to the unique property of GFP being quenched under acidic conditions [41]. Compared to the control cells, knockdown of *USP20* resulted in a reduction in autophagy flux, as evidenced by an increase in the number of autophagosomes (yellow puncta) and decrease in the number of autolysosomes (red puncta) when cultured in EBSS (**Fig. 1A**, quantification in **1B-D**). Consistently, under starvation conditions, the size of autophagosomes (yellow puncta) was increased in *USP20* knockdown cells compared to the control (**Fig. 1E**). Furthermore, when cells with *USP20* knockdown were cultured in EBSS supplemented with the lysosomal inhibitor Bafilomycin (BafA1), a mild but not significant decrease in the number of autophagosomes and an increase in their size were observed, compared to the control (**Fig. 1A**, quantification in **1B and E**). These findings strongly indicate that knockdown of *USP20* inhibits autophagy flux. Furthermore, the CRISPR-Cas9-mediated knockout of *USP20* resulted in a significant reduction in the number of LC3B puncta and the level of LC3B-II compared to the control, both under basal and starvation conditions (**Fig. 1F-H**). These results suggest that USP20 has the potential to influence both autophagy flux and autophagy initiation.

**Figure 1.**
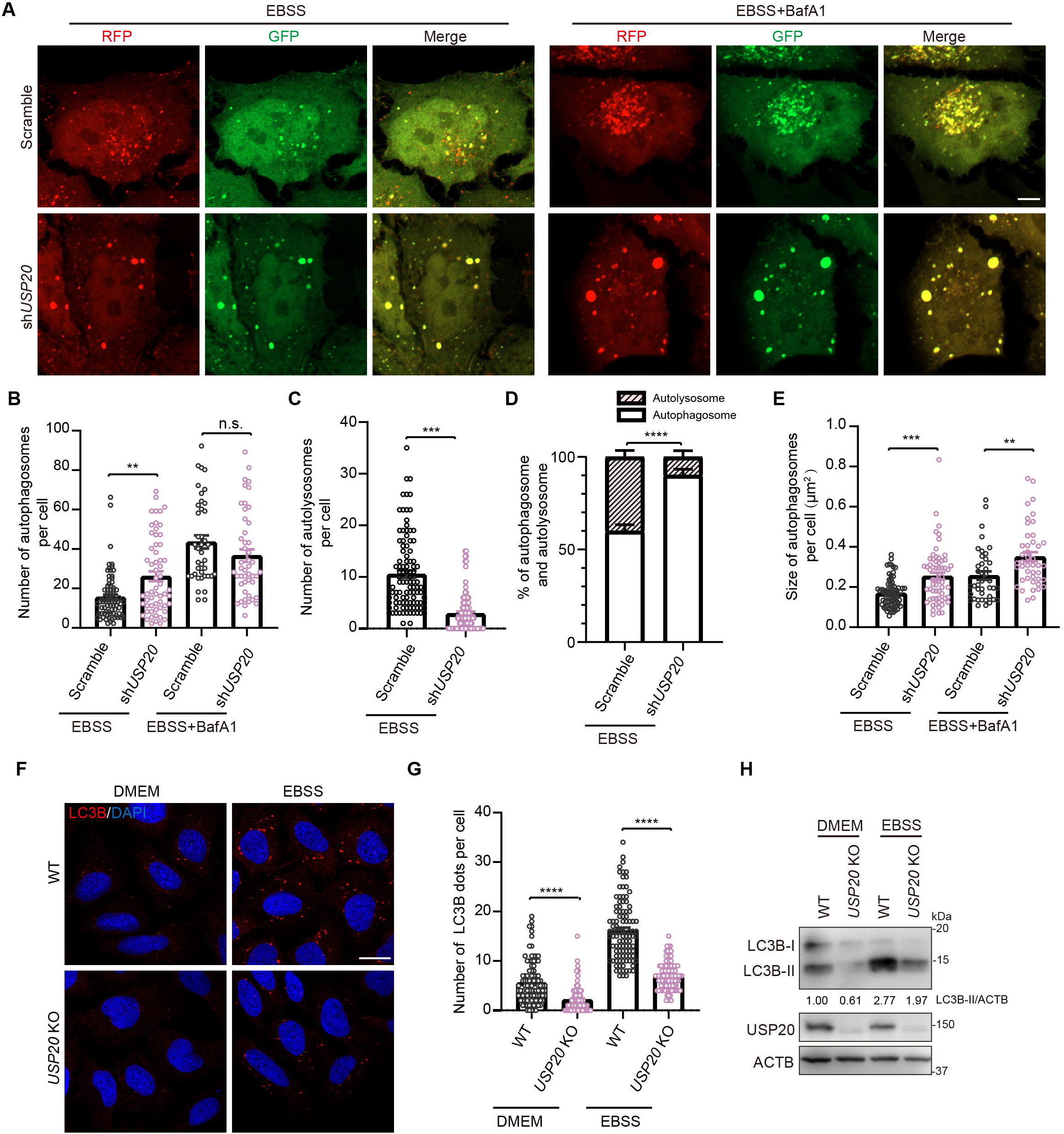
Deletion of *USP20* impairs the autophagy pathway. (**A**) HeLa cells stably expressing *RFP-GFP-LC3B* were transfected with either scramble shRNA or *USP20* shRNA. After 72 h of transfection, cells were treated with EBSS or EBSS+BafA1 (100 nM) for 4 h. Scale bar: 10 μm. (**B, C**) The number of autophagosomes (GFP^+^RFP^+^, yellow puncta) (**B**) or autolysosome (RFP^+^GFP^-^, red puncta) (**C**) per cell from (A) was quantified. (**D**) The percentage of autophagosomes (yellow) and autolysosomes (red) in a cell under starved condition using EBSS medium from (A) was determined. (**E**) The size of autophagosomes (GFP+RFP+, yellow puncta) in (A) was quantified. (**F**) *USP20* knockout inhibits autophagy. U-2 OS wild-type and *USP20* knockout cells were subjected to DMEM or EBSS treatment for 4 h prior to fixation. Subsequently, cells were immunostained with LC3B (red) and the nuclei were labeled with DAPI (blue). Scale bar: 10 μm. (**G**) The number of LC3B puncta per cell from (F) was quantified. (**H**) Immunoblotting was conducted to assess the knockdown efficiency of *USP20* and to analyze the ratio of LC3B-II to ACTB under normal and starvation conditions. ACTB was used as a loading control. Statistical analyses were performed on data from three independent experiments, with counts of more than 100 cells. Error bars represent SEM. The significance levels are indicated as n.s., not significant, \*\**p* < 0.01, and \*\*\**p* < 0.001 (one-way ANOVA with Tukey’s test).

To investigate the specific stage of the autophagy process in which USP20 is involved, we analyzed multiple complexes that have functional roles in autophagy regulation. These complexes include the ULK1 complex and the PI3K complex involved in autophagy initiation, the LC3 lipidation-related enzymes playing roles in autophagosome formation, specific ER-localized proteins that have been shown to facilitate autophagy initiation, and the SNARE complex mediating the fusion of autophagosome with lysosome. By employing immunoprecipitation to identify USP20 interacting proteins, we discovered that USP20 interacts with various proteins involved in different stages of the autophagy process. It interacts with ATG101 in the ULK1 complex, ATG14 and PIK3C3/Vps34 in the PI3K complex, VAPA, VAPB, and VMP1 on the ER membrane responsible for autophagosome membrane initiation, ATG4B in the enzyme cascade for LC3 lipidation, STX17 and VAMP8 involved in autophagosome and lysosome fusion, as well as the well-known autophagy receptor SQSTM1/p62 (**Fig. S2A-I**). However, under our experimental conditions, we did not detect its interaction with ULK1, BECN1, WIPI2, or ZFYVE1/DFCP1 (**Fig. S2I**). These findings suggest that USP20 may have an extensive role in the regulation of the autophagy pathway.

### USP20 participates in the ER-phagy process

Consistent with previous reports [42], our fluorescence imaging results confirm the colocalization of FLAG-USP20 with the ER marker protein RFP-SEC61B, despite the absence of a transmembrane domain (**Fig. 2A**). This localization pattern suggests a potential association of USP20 with ER-related functions or interactions. Based on this, we further investigated whether USP20 plays a role in regulating ER-phagy, a process involved in the degradation of ER components. To investigate the role of USP20 in regulating ER-phagy, we first employed a GFP cleavage strategy. Our results revealed a significant decrease in the cleaved GFP band in cells with *USP20* knockdown, indicating the inhibition of the ER-phagy pathway upon the reduction of USP20 levels (**Fig. 2B**, quantification in **2C**). In addition, to further assess ER-phagy, we employed the ssRFP-GFP-KDEL system [23]. Under normal conditions, there was not a significant amount of red KDEL signal observed. However, when cells were subjected to prolonged treatment with EBSS, we observed a marked increase in RFP^+^GFP^-^ (red puncta) KDEL puncta, indicating successful induction of ER-phagy (**Fig. 2E**). Conversely, when *USP20* was knocked down, we observed a significant decrease in the number of RFP^+^GFP^-^ (red puncta) KDEL puncta and a reduction in the levels of cleaved RFP under starved conditions, compared to the control (**Fig. 2D**, quantification in **2E**, and **Fig. 2F**). Consistently, overexpression of *USP20* led to an increased number of RFP^+^GFP^-^ (red puncta) KDEL puncta under starvation conditions (**Fig. S3A**, quantification in **S3B**). These observations strongly support the notion that USP20 positively regulates ER-phagy.

**Figure 2.**
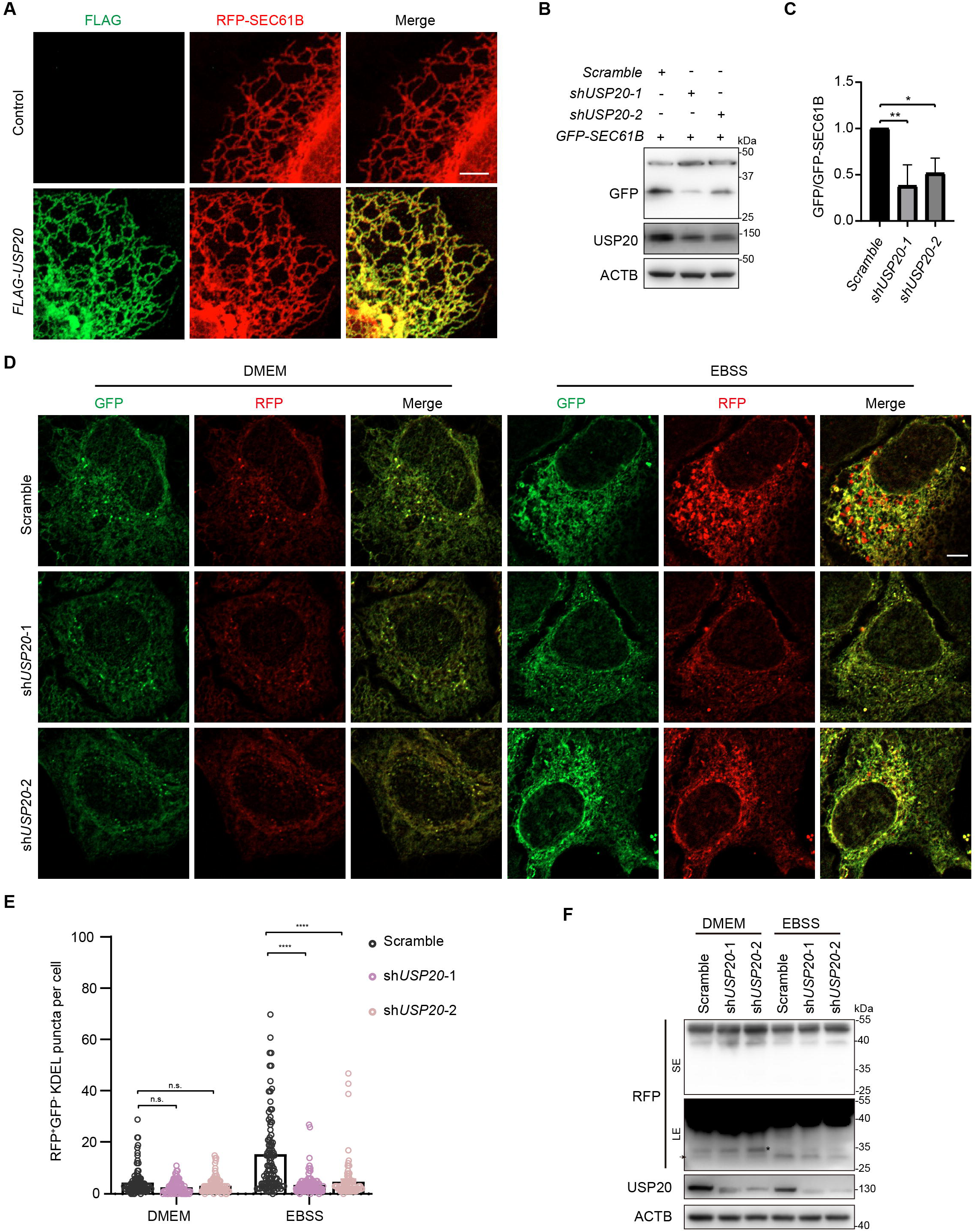
USP20 is involved in the process of ER-phagy. (**A**) U-2 OS cells were transfected with either the empty vector or *FLAG-USP20*, along with *RFP-SEC61B*. After 24 h of transfection, cells were fixed and immunostained with antibody against FLAG (green). Scale bar: 5 μm. (**B**) U-2 OS cells were transfected with scramble shRNA or *USP20* shRNA, along with *GFP-SEC61B*. After 72 h of transfection, cells were lysed with RIPA buffer, and the lysates were subjected to immunoblotting using the indicated antibodies. ACTB was used as a loading control. (**C**) The relative fold changes of the ratio of GFP to GFP-SEC61B as shown in (B) are presented as means of three replicate experiments ± SEM. The significance levels are indicated as *p < 0.05 and **p < 0.01 (one-way ANOVA with Tukey’s test). (**D**) HeLa cells stably expressing Tet-on *ssRFP-GFP-KDEL* were transfected with either scramble shRNA or *USP20* shRNA. After 48 h, doxycycline (DOX) was added to induce the expression of *ssRFP-GFP-KDEL*. After 24 h of induction, cells were treated with DMEM or EBSS for 9 h. Cells were then fixed for fluorescence detection. Scale bar: 5 μm. (**E**) The number of RFP^+^GFP^-^ (red) KDEL puncta per cell was quantified from (D). Statistical analyses were performed on data from three independent experiments, with counts of more than 100 cells. Error bars represent SEM. The significance levels are indicated as n.s., not significant, and \*\*\*\**p* < 0.0001 (one-way ANOVA with Tukey’s test). (**F**) samples from (D) were subject to immunoblotting with the indicated antibodies to assess the cleavage of ssRFP-GFP-KDEL, resulting in the generation of RFP. ACTB was used as a loading control. LE: long exposure; SE, short exposure.

### USP20 specifically mediates the deubiquitination of the ER-phagy receptor FAM134B

To investigate the regulatory mechanism of USP20 in ER-phagy, we performed a screening of currently identified ER-phagy receptors to identify the specific substrates that undergo deubiquitination by USP20. A denaturing immunoprecipitation assay was performed to examine the alterations in the ubiquitination levels of six known ER-localized ER-phagy receptors (FAM134B, ATL3, RTN3L, SEC62, CCPG1, and TEX264) upon overexpression of *USP20*. The results showed that FAM134B is specifically deubiquitinated by USP20, as the ubiquitination level of FAM134B significantly decreased under conditions of *USP20* overexpression (**Fig. S4A**). In contrast, the ubiquitination levels of the other ER-phagy receptors did not exhibit significant changes under the same conditions (**Fig. S4B-F**). This suggests that USP20 specifically targets FAM134B for deubiquitination, highlighting its selective role in regulating the ubiquitination status of this particular ER-phagy receptor.

Additionally, treatment with the specific USP20 inhibitor GSK2643943A [43] led to elevated levels of FAM134B ubiquitination compared to the control (**Fig. 3A**). Moreover, the catalytically inactive mutant of USP20, USP20^C154S/H643Q^, did not lead to a decrease in the ubiquitination level of FAM134B (**Fig. 3B**). Furthermore, the immunoprecipitation assay demonstrated that MYC-USP20 and FLAG-FAM134B exhibited an interaction, indicating their potential binding capability (**Fig. 3C**). In addition, the endogenous interaction between USP20 and FAM134B was confirmed through immunoprecipitation experiments using antibodies against FAM134B (**Fig. 3D**). Therefore, these results suggest that FAM134B is a substrate of USP20, and that the DUB activity is essential for regulating the ubiquitination of FAM134B.

**Figure 3.**
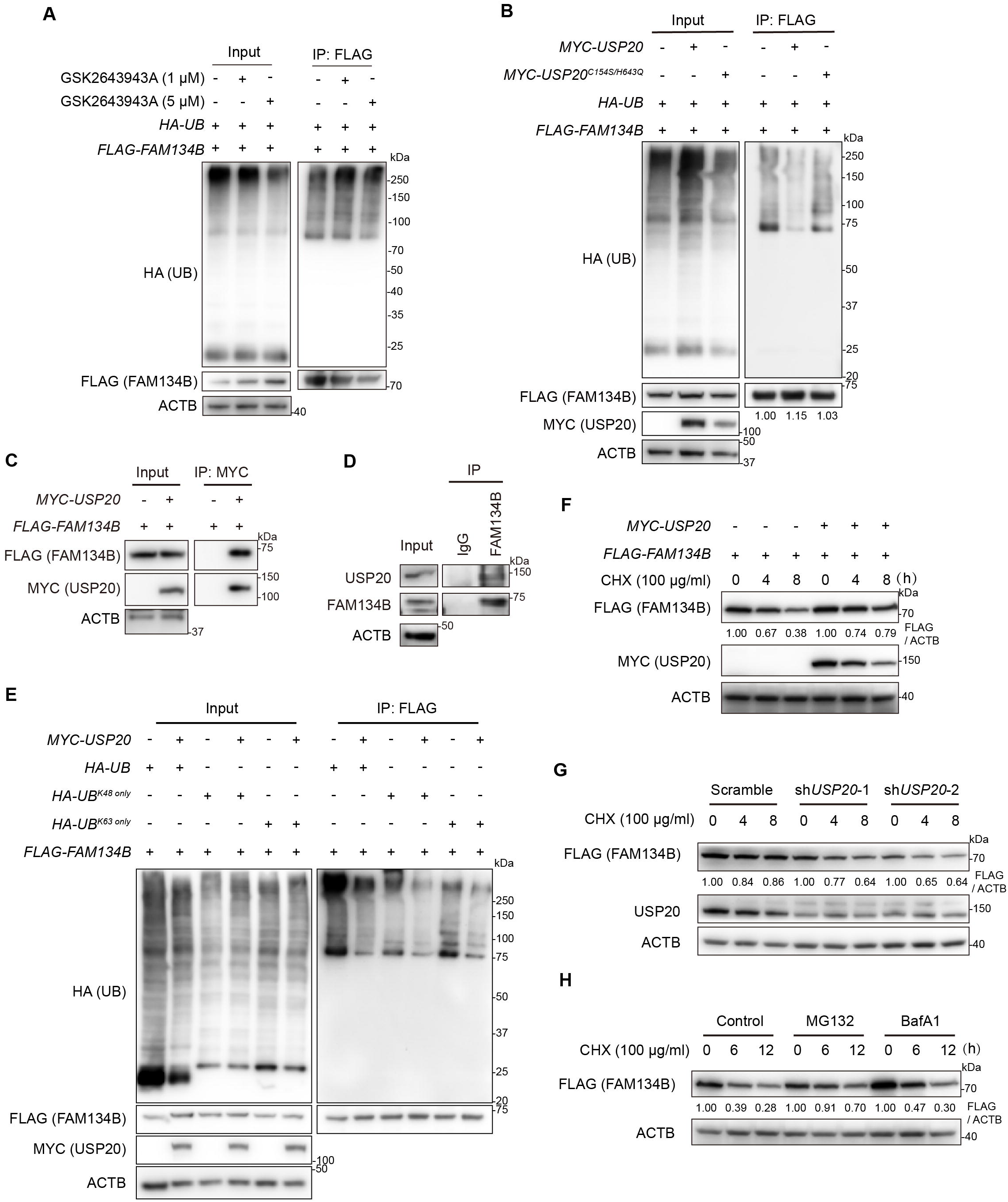
USP20 interacts with FAM134B and mediates the deubiquitination of FAM134B. (**A**) FAM134B is deubiquitinated by USP20. HEK293FT cells were transfected with *FLAG-FAM134B* and *HA-UB*. The cells were then treated with different concentration of USP20 inhibitor GSK2643943A for 12 h. Proteins were immunoprecipitated with anti-FLAG magnetic beads under denaturing conditions and immunoblotted with the indicated antibodies. (**B**) HEK293FT cells were transfected with either the empty vector, *MYC-USP20*, or *MYC-USP20^C154S/H643Q^*, along with *FLAG-FAM134B* and *HA-UB*. After 24 h of transfection, FLAG affinity isolation was performed under denaturing conditions using anti-FLAG magnetic beads, and the immunoprecipitated samples were subjected to immunoblotting analysis using the specified antibodies. The numbers indicate the relative amount of FLAG-FAM134B signal compared to the control condition. (**C**) USP20 interacts with FAM134B in cells. HEK293FT cells were transfected with either the empty vector or *MYC-USP20*, along with *FLAG-FAM134B*. MYC affinity isolation was performed using anti-MYC magnetic beads, and the samples were subjected to immunoblotting analysis using the specified antibodies. (**D**) The endogenous interaction between USP20 and FAM134B. Cell lysates from HEK293FT cells were subjected to immunoprecipitation using antibodies against FAM134B. The immunoprecipitated samples were then analyzed by immunoblotting to detect the presence of USP20. (**E**) USP20 cleaves both K48– and K63-linked ubiquitin chains from FAM134B. HEK293FT cells were transfected with either the empty vector or *MYC-USP20*, together with *HA-UB* or *HA-UB^K48^ ^only^* or *HA-UB^K63^ ^only^*, along with *FLAG-FAM134B*. FLAG affinity isolation was performed under denaturing conditions using anti-FLAG magnetic beads, and the immunoprecipitated samples were subjected to immunoblotting analysis using the specified antibodies. (**F** and **G**) Cycloheximide (CHX) chase analysis of FLAG-FAM134B under control or *USP20* overexpression conditions (**F**), or under scramble shRNA or *USP20* shRNA knockdown conditions (**G**). HEK293FT cells were treated with CHX at a concentration of 100 μg/ml for 0, 4, or 8 h to block protein synthesis. The degradation rate of FLAG-FAM134B was monitored over time by immunoblotting. (**H**) FLAG-FAM134B protein degradation is proteasome-dependent. HEK293FT cells expressing *FLAG-FAM134B* were treated with a control treatment, MG132 (10 μM), or BafA1 (100 nM) for 12 h. The cells were treated with CHX at a concentration of 100 μg/ml for 0, 6, or 12 h to block protein synthesis. The degradation rate of FLAG-FAM134B was then monitored over time by immunoblotting. ACTB was used as the loading control in all the immunoblotting analyses. The numbers indicate the relative amount of FLAG-FAM134B signal compared to the time 0 point, normalized by ACTB.

### USP20-mediated deubiquitination of FAM134B facilitates the stabilization of **FAM134B**

Previous studies have shown that USP20 preferentially hydrolyzes K48– and K63-linked ubiquitin chain and reduces the ubiquitination of HMGCR (3-hydroxy-3-methylglutaryl-coenzyme A reductase), resulting in substrate stabilization, activation, and subsequent influence on cholesterol biogenesis within the liver after feeding [43]. To investigate the deubiquitination patterns of USP20 on FAM134B, we utilized ubiquitin variants that specifically form K48 or K63 polyubiquitin linkages (referred to as UB^K48^ ^only^ and UB^K63^ ^only^, respectively) in the ubiquitination assay. Denaturing immunoprecipitation experiments confirmed that USP20 can remove both K48 and K63-linked ubiquitin chains from FAM134B (**Fig. 3E**). Subsequent investigations were conducted to explore the involvement of USP20 in the regulation of FAM134B degradation. Our findings demonstrated that overexpression of *USP20* led to a slight increase in FAM134B levels compared to the control cells, which was dependent on the DUB activity of USP20 (**Fig. 3B**). Since FAM134B is a protein with high stability [23], its degradation is not easily detectable at the steady-state level. We therefore conducted cycloheximide chase experiments to assess its degradation kinetics. Remarkably, our findings demonstrated that *USP20* overexpression significantly enhanced the stability of FAM134B, whereas *USP20* knockdown led to reduced FAM134B protein stability (**Fig. 3F**-**G**). Treatment with the proteasome inhibitor MG132, but not the lysosome inhibitor BafA1, effectively blocked the degradation of FAM134B protein, as demonstrated by the accumulation of FAM134B levels compared to untreated cells (**Fig. 3H**). These findings indicate that USP20 participates in the regulation of FAM134B degradation through the proteasome pathway.

Next, we searched for the specific sites where USP20 deubiquitinates FAM134B. FAM134B has been previously reported to undergo ubiquitination primarily at K160 and K247 [44, 45]. We therefore investigated whether mutations in these two lysine residues led to decreased ubiquitination levels and impaired cleavage by USP20. Surprisingly, we observed that mutating K160 and K247 did not result in a significant reduction in FAM134B ubiquitination levels (**Fig. S5A-B**). Furthermore, USP20 was still able to effectively deubiquitinate FAM134B even in the presence of these mutations. These findings suggest that there may be other lysine residues or alternative modifications that contribute to the ubiquitination and subsequent regulation of FAM134B.

The FAM134B protein sequence contains 24 lysine residues, distributing in the transmembrane RHD domain and the C-terminus (**Fig. S5A**). We next generated a series of FAM134B truncation mutants, specifically deleting either the RHD or the C-terminal domain (**Fig. S5C**). The mapping results showed that USP20 was capable of interacting with all the truncation mutants of FAM134B (**Fig. S5D**). Additionally, denaturing immunoprecipitation experiments provided evidence that USP20 could still deubiquitinate FAM134B when its RHD domain or the C-terminal domain was deleted (**Fig. S5E**). These results suggest that FAM134B has multiple ubiquitination sites, and USP20 exerts comprehensive deubiquitination regulation on FAM134B.

### USP20 promotes autophagy by facilitating the interaction between FAM134B and LC3B

FAM134B serves as an essential receptor for ER-phagy and plays a critical role in the process by interacting with LC3. Next, we sought to determine whether the increased expression of *USP20* affects the interaction between FAM134B and LC3B. Our results demonstrated a significant increase in the number of LC3B puncta colocalized with FAM134B in *USP20*-overexpressing cells compared to control cells under normal conditions and EBSS plus BafA1 treatment (**Fig. 4A**, quantification in **4B**). In addition, our immunoprecipitation results demonstrate that the interaction between FAM134B and LC3B is notably strengthened upon overexpression of *USP20* (**Fig. 4C**). This suggests that USP20 may positively regulate the association between FAM134B and LC3, leading to the promotion of ER-phagy.

**Figure 4.**
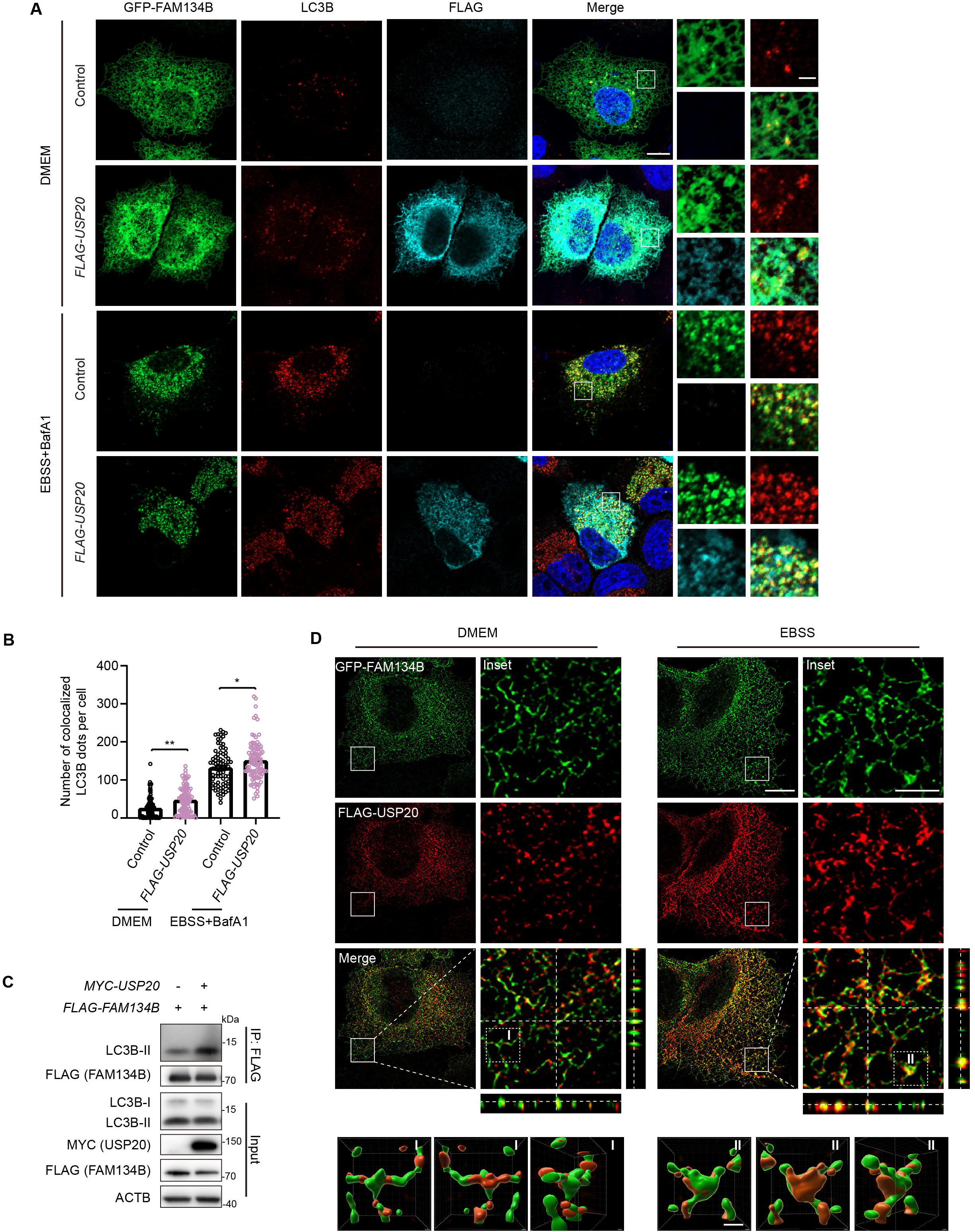
USP20 promotes the interaction between FAM134B and LC3B. (**A**) HeLa cells were transfected with either the empty vector or *FLAG-USP20*, along with *GFP-FAM134B*. Following transfection, the cells were treated with either DMEM or EBSS+BafA1 (100 nM) for 9 h before fixation. Subsequently, cells were immunostained with FLAG (cyan) and LC3B (red), and nuclei were labeled with DAPI (blue). Scale bar: 10 μm. The insets indicate the enlarged area. Scale bar: 2 μm. (**B**) The number of colocalized LC3B with FAM134B per cell was quantified. Statistical analyses were performed on data from three independent experiments, with counts of more than 100 cells. Error bars represent SEM. The significance levels are indicated as *p < 0.05 and \*\**p* < 0.01 (one-way ANOVA with Tukey’s test). (**C**) FAM134B interacts with a higher amount of LC3B upon *USP20* overexpression. HEK293FT cells were co-transfected with either the control vector or *FLAG-FAM134B*, along with the vector or *MYC-USP20.* Immunoprecipitation was performed using anti-FLAG magnetic beads, and the samples were immunoblotted with the indicated antibody. ACTB was used as a loading control. (**D**) Increased colocalization of USP20 with FAM134B during starvation. HeLa cells were transfected with *FLAG-USP20* and *GFP-FAM134B*. After transfection, the cells were treated with either DMEM or EBSS for 9 h before fixation. Immunostaining was performed using antibodies against FLAG (red). Z-stack projection of representative images showing the signals of FLAG-USP20 and GFP-FAM134B was acquired by SIM. Scale bar: 10 μm. The insets indicate magnified orthogonal sectioning views of regions within the boxes. Scale bar: 2 μm. The area within Box I and II is rendered in 3D and displayed from different angles. Scale bar: 0.5 μm.

To gain further insights into the localization pattern of FAM134B and USP20, we employed super-resolution structured illumination microscopy (SIM) to resolve their distribution on the ER. Our observations revealed that both FAM134B and USP20 exhibit a non-uniform distribution on the ER membrane (**Fig. 4D**). Under normal conditions, USP20 was found to colocalize with FAM134B specifically in areas where FAM134B formed punctate structures (**Fig. 4D**). Remarkably, under starved conditions, we observed an increased colocalization between USP20 and FAM134B, accompanied by a greater number of FAM134B puncta (**Fig. 4D**). These findings suggest that USP20 plays a role in stabilizing FAM134B and facilitating the recruitment of LC3 to FAM134B, thereby promoting ER-phagy in response to nutrient deprivation.

### VAPs mediate the localization of USP20 to the ER

Since USP20 lacks a transmembrane domain but localizes to the ER, we were intrigued by whether it is anchored to the ER by specific ER proteins. Remarkably, USP20 contains two FFAT motifs, which are typical motifs known to interact with ER-transmembrane proteins VAPA/B (**Fig. 5A**) [46]. To gain insight into this mechanism, we conducted investigations into the interaction between USP20 and VAPs. VAPs have been previously reported to interact with proteins that contain FFAT motifs via their MSP domain, resulting in their localization to the ER [47]. Immunoprecipitation experiments provided evidence that USP20 FFAT1, along with FFAT2, plays a central role in mediating the interaction between USP20 and VAPs (**Fig. 5B**). To further verify the FFAT-dependent localization of USP20 to the ER, we conducted a microsome fractionation experiment. The results demonstrated that wild-type USP20 predominantly localized to the microsomes derived from the ER, whereas the FFAT1/2-deleted mutant of USP20 was primarily found in the cytosol (**Fig. 5C**). Furthermore, the immunofluorescence imaging analysis demonstrated that upon loss of the FFAT1/2 motifs, USP20 exhibited a cytoplasmic localization (**Fig. 5D**). The VAPA^K94D/M96D^ and VAPB^K87D/M89D^, mutants that lost the interaction with FFAT motif [12], exhibited a reduced interaction with USP20 compared to the wild-type VAPA/B (**Fig. 5E**). In addition, USP20 exhibits a higher distribution in the cytosol fraction in VAPA/B double knockout cells compared to the control cells (**Fig. 5F**). Based on these results, we concluded that VAPs mediate the ER localization of USP20.

**Figure 5.**
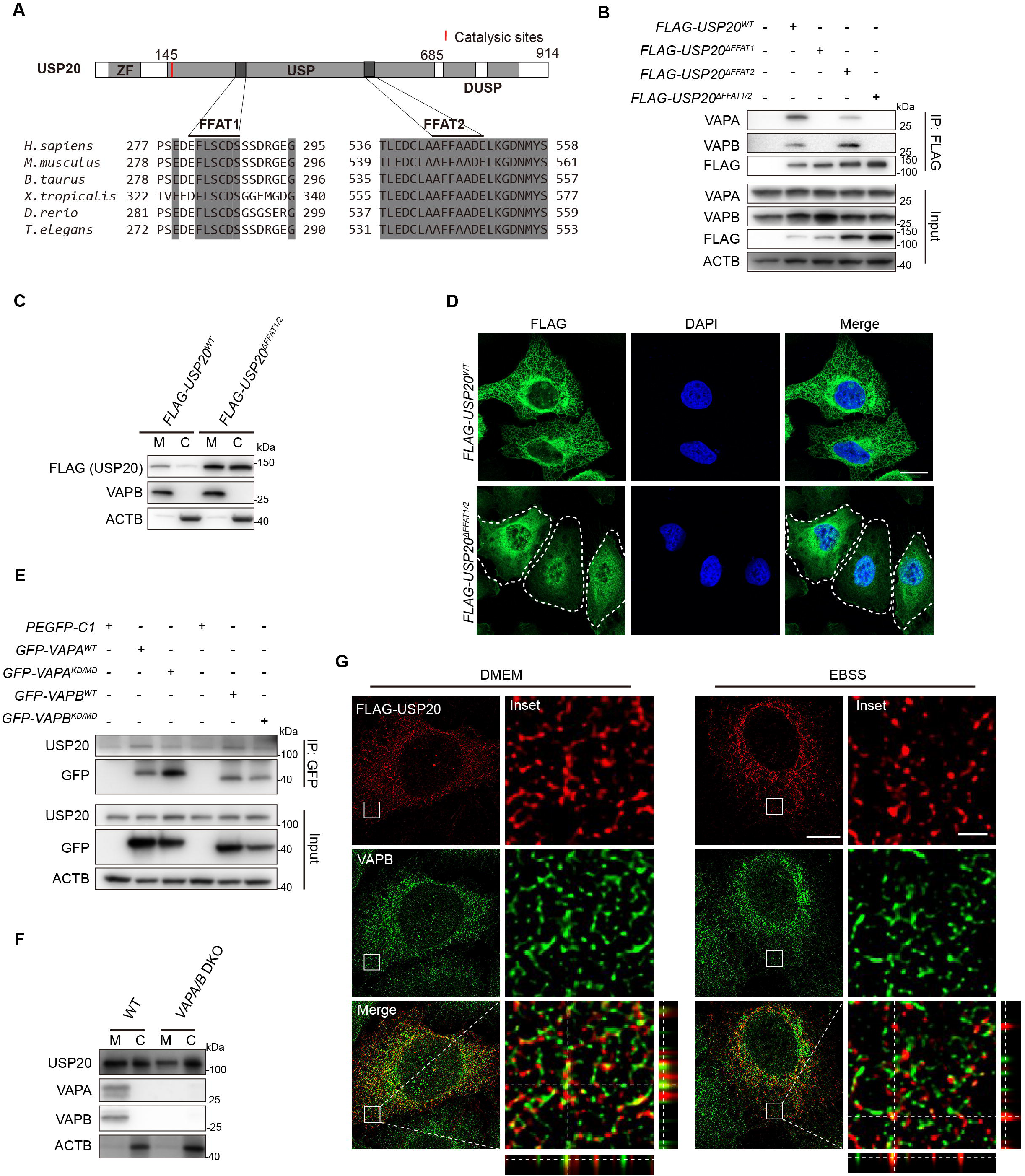
VAPs mediate the localization of USP20 to the ER. (**A**) The protein sequence of USP20 contains multiple domains, including a zinc finger (ZF) domain, a catalytic USP domain, and two DUSP domains. Two FFAT motifs located in the USP domain are conserved among different organisms. Multiple sequence alignment of USP20 proteins from various species was performed using the Clustal Omega in the European Bioinformatics Institute (EMBL-EBI) (https://www.ebi.ac.uk/Tools/msa/clustalo/). Conserved regions are shaded in grey. (**B**) The FFAT motifs of USP20 are responsible for its interaction with VAPA/B proteins. HEK293FT were transfected with *FLAG-USP20* wild-type, or *FLAG-USP20*^Δ*FFAT1*^ or *FLAG-USP20*^Δ*FFAT2*^ or *FLAG-USP20*^Δ*FFAT1/2*^ mutants. FLAG affinity isolation was performed using anti-FLAG magnetic beads, and the samples were analyzed by immunoblotting with the indicated antibodies. ACTB was used as a loading control. (**C**) Mutation of the FFAT motifs of USP20 results in a reduction in ER enrichment. HEK293FT cells were transfected with either *FLAG-USP20* wild-type or *FLAG-USP20*^Δ*FFAT1/2*^ mutant. A microsome fractionation assay was performed to demonstrate the enrichment of USP20 in the ER. The ER marker VAPB was used to indicate the ER fraction, while ACTB was used to indicate the cytosol fraction. (**D**) FLAG-USP20^ΔFFAT1/2^ mutant does not localize to the ER. HeLa cells were transfected with either *FLAG-USP20* wild-type or *FLAG-USP20*^Δ*FFAT1/2*^ mutant. After 24 h of transfection, the cells were fixed and immunostained with anti-FLAG antibody (green) and the nuclei were labeled with DAPI (blue). Scale bar: 10 μm. (**E**) VAPA/B proteins interact with less USP20 when their FFAT-binding motifs are mutated. HeLa cells were transfected with either *GFP-VAPA/B* wild-type or *GFP-VAPA^KD/MD^* or *GFP-VAPB^KD/MD^* mutants. After 24 h of transfection, GFP immunoprecipitation assay was performed, and the samples were analyzed by immunoblotting using the specified antibodies. ACTB was used as a loading control. (**F**) USP20 shows reduced distribution in the ER when *VAPA/B* are knocked out. A microsome fractionation assay was performed using both HEK293FT wild-type cells and *VAPA/B* double knockout cells to evaluate the enrichment of USP20 in the ER. The efficiency of VAPA/B knockdown was confirmed by immunoblotting with VAPA and VAPB antibodies, while ACTB was used as a marker for the cytosol fraction. (**G**) USP20 and VAPB localization on ER. HeLa cells were transfected with *FLAG-USP20*. Following transfection, the cells were subjected to DMEM or EBSS treatment for 9 h prior to fixation. Subsequently, cells were immunostained with FLAG (red) and VAPB (green). Z-stack projection of representative images showing the signals of FLAG-USP20 and VAPB was acquired by SIM. Scale bar: 10 μm. The insets indicate magnified orthogonal sectioning views of regions within the boxes. Scale bar: 1 μm.

We then used SIM to investigate the localization of USP20 and VAPB on the ER membrane. The two proteins were not evenly distributed on the ER membrane and exhibited an intercalated localization pattern with overlapping signals to some extent under normal conditions (**Fig. 5G**). Following treatment with EBSS, we observed a specific colocalization of USP20 and VAPB at the sites where USP20 forms puncta (**Fig. 5G**). These findings suggest that USP20 and VAPB are closely localized within the ER. Under starvation conditions, both VAPB and USP20 undergo transfer to specific areas, providing further evidence for their potential functional interaction in the regulation of ER-phagy.

### USP20 facilitates FAM134B-VAPB association and enhances early autophagy protein recruitment for ER-phagy

We utilized SIM to study the localization of FAM134B and VAPB on the ER membrane, revealing an adjacent localization pattern with partial overlapping signals under normal conditions (**Fig. 6A**). Upon EBSS treatment, a specific colocalization of FAM134B and VAPB was observed at the sites where FAM134B forms puncta (**Fig. 6A**). We then investigated the potential involvement of VAPA/B in mediating the deubiquitination of FAM134B by USP20. Surprisingly, even in the presence of the USP20 FFAT mutant, we observed that USP20 was still capable of binding and deubiquitinating FAM134B (**Fig. S6A-B**). These findings suggest that while VAPA/B may play a role in mediating the localization of USP20 to the ER, they may have additional functions beyond facilitating USP20-mediated deubiquitination of FAM134B.

**Figure 6.**
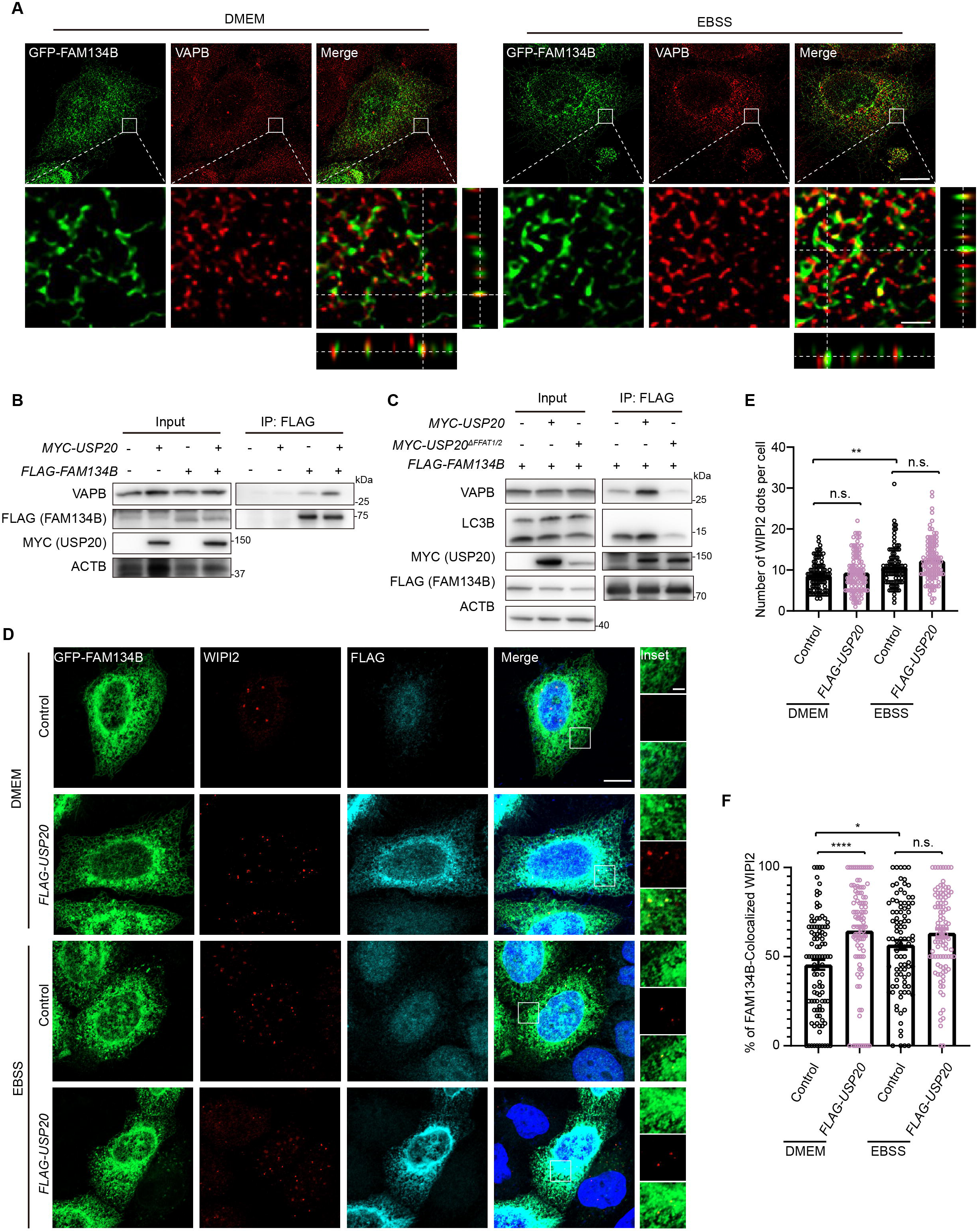
USP20 enhances the interaction between FAM134B and VAPB on the ER. (**A**) FAM134B and VAPB localization on ER. HeLa cells were transfected with *GFP-FAM134B*. Following transfection, the cells were subjected to DMEM or EBSS treatment for 9 h prior to fixation. Subsequently, cells were immunostained with VAPB (red). Z-stack projection of representative images showing the signals of GFP-FAM134B and VAPB was acquired by SIM. Scale bar: 10 μm. The insets indicate magnified orthogonal sectioning views of regions within the boxes. Scale bar: 2 μm. (**B**) USP20 promotes FAM134B interaction with VAPB. HEK293FT cells were co-transfected with either the empty vector or *MYC-USP20*, along with the empty vector or *FLAG-FAM134B*. FLAG affinity isolation was performed using anti-FLAG magnetic beads, and the immunoprecipitated samples were subjected to immunoblotting analysis using the specified antibodies. ACTB was used as a loading control. (**C**) The USP20 FFAT mutant does not enhance the interaction between FAM134B and VAPB. HEK293FT cells were co-transfected with either the empty vector or *MYC-USP20* or *MYC-USP20*^Δ*FFAT1/2*^, along with *FLAG-FAM134B*. FLAG affinity isolation was performed using anti-FLAG magnetic beads, and the immunoprecipitated samples were subjected to immunoblotting analysis using the specified antibodies. ACTB was used as a loading control. (**D**) Overexpression of *USP20* leads to a significant increase in the colocalization of WIPI2 with FAM134B under normal conditions and a marginal increase under starvation conditions. HeLa cells were transfected with the control vector or *FLAG-USP20*, along with *GFP-FAM134B*. Following transfection, the cells were subjected to DMEM or EBSS treatment for 9 h prior to fixation. Subsequently, cells were immunostained with WIPI2 (red) and FLAG (cyan). Scale bar: 10 μm. The insets indicate the enlarged area. Scale bar: 2 μm. (E and F) The number of WIPI2 dots per cell (E) and the percentage of FAM134B-colocalized WIPI2 (F) from (A) was quantified.

Previous studies have demonstrated that VAPs promote autophagy by facilitating ER/isolation membrane contact through binding with WIPI2 and recruiting various autophagy proteins to facilitate the assembly of the autophagy initiation complex at the ER [12, 48]. In our study, we observed an enhanced interaction between FAM134B and VAPB upon overexpression of *USP20* (**Fig. 6B**). Conversely, their interaction was significantly reduced when expressing the *USP20* FFAT mutant compared to wild-type *USP20* (**Fig. 6C**). Consistently, we observed that FAM134B exhibited reduced binding to LC3 proteins when USP20 was unable to localize to the ER (**Fig. 6C**). Additionally, we observed a significant increase in the colocalization of FAM134B with the early autophagy protein WIPI2 upon *USP20* overexpression under normal conditions. Although there was a mild increase in colocalization under starvation conditions, the p value did not reach significance (**Fig. 6D**, quantification in **6E-F**). These findings suggest that USP20 may enhance the association between FAM134B and VAPB, thereby playing a role in facilitating ER-phagy by recruiting more early autophagy proteins to the initiation sites.

## Discussion

Our study identified USP20 deubiquitinates and stabilizes the ER-phagy receptor FAM134B to facilitate ER-phagy. USP20 has previously been shown to regulate autophagy by deubiquitinating and stabilizing ULK1, thus promoting autophagy [36]. Given the localization of USP20 to the ER, it prompted us to investigate whether USP20 regulates the deubiquitination of ER-targeted proteins. In this study, we elucidated the recruitment of USP20 to the ER by VAP proteins, its role in deubiquitinating the ER-phagy receptor FAM134B, and its function in stabilizing FAM134B to promote ER-phagy (**Fig. 7**).

**Figure 7.**
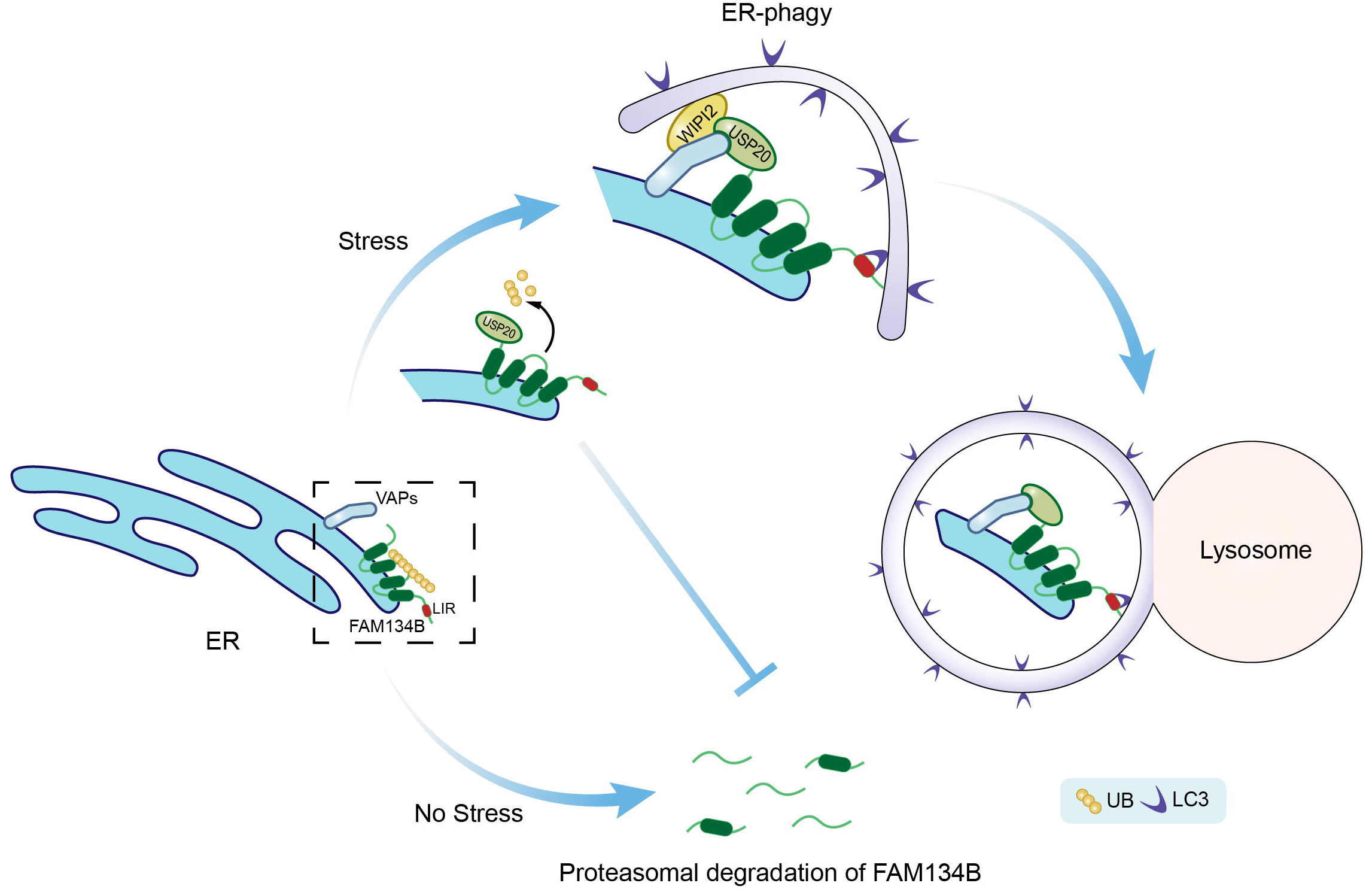
A proposed model for the role of USP20 in regulating FAM134B deubiquitination and facilitating ER-phagy. USP20 deubiquitinates and stabilizes FAM134B, while also being recruited to the ER by VAPs. Under starvation conditions, USP20 and FAM134B form puncta. Additionally, USP20 recruits VAPs to the region where both USP20 and FAM134B are enriched. This further leads to the recruitment of early autophagy proteins including WIPI2, thereby facilitating ER-phagy.

Previous studies on USP20 have focused on its activity in metabolic diseases, antiviral immunity and cancer pathogenesis [43, 49, 50]. For example, the phosphorylated USP20 binds to ER-localized AMFR/gp78 and stabilizes HMGCR through cleavage of K48– and K63-linked ubiquitin chains, leading to an increase in cholesterol biosynthesis within the liver after feeding [43]. Furthermore, USP20 plays a pivotal role in cellular antiviral responses by promoting deconjugation of K48-linked ubiquitin chains from STING1/MITA, leading to enhanced stability of STING1 and increased expression of type I IFNs and proinflammatory cytokines following DNA virus infection [49, 50]. USP20 reduces atherogenic signaling in smooth muscle cells by deubiquitinating RIPK1 in response to TNF and IL1B stimulation [51]. Moreover, study reported that USP20 promotes breast cancer metastasis by stabilizing the metastasis-promoting transcription factor SNAI2 [52]. The research findings also revealed a correlation between higher expression of USP20 and worse metastasis-free survival, which is consistent with the observations made in mice, highlighting the role of USP20 in promoting metastasis [52]. Additional investigations have unveiled USP20’s involvement in the deubiquitination and stabilization of diverse substrates, including ADRB2 and SQSTM1 [53, 54]. Notably, USP20 collaborates with USP33 in the efficient removal of ubiquitin chains from tail-anchored proteins, following their insertion into the ER. This process rescues posttranslationally targeted membrane proteins that have been inappropriately ubiquitinated by protein quality control in the cytosol, underscoring the indispensable role of USP20 in protein homeostasis [42]. In the present study, we reveal the novel role of USP20 in the cleavage of K48 and K63-linked ubiquitin chains on FAM134B, ultimately leading to the stabilization of FAM134B and facilitating the process of ER-phagy. Thus, our findings provide compelling evidence for USP20’s multifaceted engagement in diverse pathways, including ER-phagy, where it exerts its deubiquitinating and substrate-stabilizing effects.

USP20 lacks a predicted transmembrane domain, yet it localizes to the ER. Our investigation here identifies the pivotal role of the FFAT domains in mediating USP20’s ER localization. Our experiments revealed that mutations in the FFAT domains of USP20 disrupt its localization to the ER, underscoring the critical importance of these domains in this process (**Fig. 5A-F**). Surprisingly, even with the FFAT mutations, USP20 retains its ability to interact with ER-localized FAM134B and carry out deubiquitination of FAM134B (**Fig. S6A-B**). The mechanism underlying this unique binding and deubiquitination ability of the USP20 FFAT mutant to ER-localized FAM134B remains yet to be elucidated. Furthermore, our investigation has shed light on the interplay between the FFAT domains of USP20 and VAPs, which serves as a key mechanism for recruiting VAPs to specific ER subdomains (**Fig. 5G**). Through this orchestrated recruitment, USP20 facilitates the subsequent assembly of autophagy initiation proteins at these sites, culminating in the promotion of efficient ER-phagy (**Fig. 6D-F**). This intricate interplay between USP20, VAPs, and the autophagy machinery emerges as a crucial determinant in the maintenance of ER homeostasis, unraveling novel insights into the regulation of ER-phagy in response to cellular stress.

ER-phagy relies on ER-localized receptors that typically present LIR motifs to interact with LC3 and initiate the autophagic process. However, the mechanism of how the phagophore is initiated and extended during ER-phagy has not been previously elucidated. Previous studies have demonstrated that certain ER proteins engage in interactions with autophagy initiation proteins, contributing to the regulation of autophagosome biogenesis [12, 48]. In the current study, we provide evidence that ER-phagy also requires a membrane recruitment event, which subsequently provides lipidated-LC3 for interaction with the ER-phagy receptor FAM134B. This interaction is essential for the efficient execution of ER-phagy.

In summary, our findings demonstrate that USP20 plays a crucial role in deubiquitinating and stabilizing the ER-phagy receptor FAM134B at specific ER subdomains. Additionally, USP20’s interaction with VAPs further enhances its role in promoting efficient ER-phagy.

## Materials and Methods

### Cell culture

HEK293FT (PTA-5077), HeLa (CRM-CCL-2) and U-2 OS (HTB-96) cells obtained from ATCC were cultured in DMEM (Thermo Fisher Scientific, 12800017) supplemented with 10% fetal bovine serum (Lonsera, S711-001S) and 100 U/ml penicillin G and 100 μg/ml streptomycin (Thermo Fisher Scientific, 15140148) at 37°C under 5% CO_2_. For autophagy and ER-phagy induction experiments, cells were subject to Earle’s Balanced Salt Solution (EBSS) (Sigma-Aldrich, E2888) for 4 h and 9 h, respectively.

### Generating knockout cell lines

The knockout cell lines were generated using the CRISPR-Cas9 system. Guide RNAs sequences were designed using the website http://chopchop.cbu.uib.no/. The guide RNAs sequences were inserted into the *pGL3-U6-sgRNA-PGK-puromycin* plasmid (Addgene 51133, Xingxu Huang). U-2 OS, HeLa or HEK293FT cells were co-transfected with the sgRNA plasmid and the *pST1374-NLS-flag-linker-Cas9* (Addgene 44758, Xingxu Huang). After 24 h, cells were selected with 5 μg/ml puromycin (Sigma-Aldrich, 540411) and 10 μg/ml blasticidin (Thermo Fisher Scientific, R21001) for 7 days. The knockout of the target gene was confirmed by immunoblotting. Single-cell colonies were then sorted by flow cytometry into 96-well plates, and each colony was screened by immunoblotting. The target sequences used are listed in **Table S1**.

### Constructing inducible stable cell lines

To generate lentiviral particles, HEK293FT cells were co-transfected with the Tet-on *ssRFP-GFP-KDEL* plasmid (Addgene 128257, Noboru Mizushima) and the lentivirus packaging plasmids *psPAX2* (Addgene 12260, Didier Trono) and *pMD2.G* (Addgene 12259, Didier Trono) using Lipofectamine® 2000 (Thermo Fisher Scientific, 11668019). After 48 h of transfection, lentiviral particles were harvested and centrifuged to remove cell debris. Hela cells were then plated in 6-well plates and infected with the viral particles for 24 h. Subsequently, the virus particle-containing medium were replaced with fresh medium. After 48 h of infection, cells were selected with 2 μg/ml puromycin. Doxycycline treatment was applied for 24 h to induce the expression of *ssRFP-GFP-KDEL*.

### Plasmids

*FLAG-USP20* and *FLAG-FAM134B* constructs were generated by cloning the corresponding cDNA into the *pRK5-FLAG* vector [55]. *MYC-USP20* and *MYC-FAM134B* constructs were generated by inserting the corresponding cDNA into the *pRK5-MYC* vector. *FLAG-USP20*^Δ*FFAT1*^*, FLAG-USP20*^Δ*FFAT2*^*, FLAG-USP20*^Δ*FFAT1/2*^ and *MYC-USP20^C154S/H643Q^* constructs were generated by performing overlap extension of PCR-mediated deletions on *pRK5-FLAG-USP20* and *pRK5-MYC-USP20*. *FLAG-FAM134B*^Δ*RHD*^*, FLAG-FAM134B*^Δ*C*^*, MYC-FAM134B*^Δ*RHD*^ and *MYC-FAM134B*^Δ*C*^ constructs were generated by overlap extension of PCR-mediated deletions on *pRK5-FLAG-FAM134B* and *pRK5-MYC-FAM134B. EGFP-SEC61B* was generated by cloning the corresponding cDNA into the *pEGFP.C1* vector (Clontech, 6084-1). The plasmid *pRK5-HA-UB* was described previously [56]. *HA-UB^K48only^* and *HA-UB^K63only^* were generated through PCR-mediated site-directed mutagenesis on *pRK5-HA-UB.* All mutations were confirmed by DNA sequencing. The plasmids encoding *GFP-VAPA, GFP-VAPA^KD/MD^*, *GFP-VAPB, GFP-VAPB^KD/MD,^ GFP-VMP1, GFP-STX17* and *GFP-VAMP8* were kindly provided by Dr. Hong Zhang (Institute of Biophysics, Chinese Academy of Sciences).

### Antibodies and reagents

Mouse anti-MYC (2276), rabbit anti-HA (3724) and rabbit anti-FAM134B (83414S, for immunoprecipitation) were purchased from Cell Signaling Technology. Rabbit anti-FLAG (F3165), mouse anti-FLAG (F7425) and rabbit anti-MAP1LC3B (L7543) were purchased from Sigma-Aldrich. Rabbit anti-FAM134B (21537-1-AP, for immunoblotting), rabbit anti-VAPA (15275-1-AP) and rabbit anti-VAPB (14477-1-AP) were purchased from Proteintech. Rabbit anti-LC3B (A19665, for immunoblotting), rabbit anti-SQSTM1 (A11247), rabbit anti-MYC (AE070) and mouse anti-HA (AE008) were purchased from ABclonal, Inc. Mouse anti-MYC (sc-166247) was purchased from Santa Cruz Biotechnology. Rabbit anti-WIPI2 (ab105459) was purchased from Abcam. Rabbit anti-USP20/VDU2 (A301-189A) was purchased from Bethyl Laboratories. The rabbit polyclonal antibody against GFP was generated using a recombinant GFP protein derived from *Aequorea victoria*. Mouse anti-LC3B (M152-3, for immunofluorescence), Anti-ACTB (HRP-DirecT; PM053-7) were purchased from Medical & Biological Laboratories Co., LTD. The secondary antibodies goat anti-mouse IgG (H+L), HRP (111-035-146), goat anti-rabbit IgG (H+L), HRP (111-035-144), and Cy5-AffiniPure donkey anti-rabbit IgG (H+L) (711-175-152) were purchased from Jackson ImmunoResearch Inc. The secondary antibodies goat anti-rabbit IgG (H+L), Alexa Fluor 488 (A-11034), goat anti-mouse IgG (H+L), Alexa Fluor 488 (A-11029), goat anti-rabbit IgG (H+L), Alexa Fluor 568 (A-11036), goat anti-mouse IgG (H+L), Alexa Fluor 568 (A-11031), and goat anti-mouse IgG (H+L), Alexa Fluor 633 (A-21052) were purchased from Invitrogen. MG132 (S2619) and cycloheximide (S7418) were purchased from Selleck. Chloroquine (C6628) was purchased from Sigma-Aldrich. Bafilomycin A1 (sc-201550A) was purchased from Santa Cruz Biotechnology. Doxycycline (60204ES03) was purchased from Yeasen. GSK2643943A (HY-111458) was purchased from MedChemExpress (MCE).

### Immunoprecipitation and immunoblotting

For immunoprecipitation, HEK293FT cells were transfected and lysed 24 h post-transfection in NP40 lysis buffer (50 mM Tris-HCl, pH 7.5, 150 mM NaCl, 5 mM MgCl_2_, 0.5% NP40 [Sangon Biotech, A600385-0100]) containing EDTA-free protease inhibitor cocktail (Bimake, B14002) for 30 min at 4°C. The lysates were centrifuged at 17,000 x g for 10 min at 4°C, and the soluble supernatant fractions were subjected to immunoprecipitation. Immunoprecipitation was performed using protein A beads (Bimake, B23202) pre-bound with FAM134B or GFP antibody, FLAG (Bimake, B26102), HA (Bimake, B26201), or MYC (Bimake, B26302) magnetic beads. After incubation for 2 h at 4°C, the beads were washed three times with NP40 wash buffer (50 mM Tris-HCl, pH 7.5, 150 mM NaCl, 5 mM MgCl_2_, 0.1% NP40), and the samples were immunoblotted with the indicated antibodies.

For immunoprecipitation under denaturing condition, harvested cells were lysed in a buffer with 1% SDS (Sinopharm Chemical Reagent Co., 30166480) and 5 mM DTT (MDBio, Inc., D023) in PBS (Sangon Biotech, B548117-0500). The samples were heated at 65°C for 15 min and diluted with NP40 lysis buffer. The soluble supernatant fractions were harvested and subjected to immunoprecipitation as described above.

Immunoblotting was performed using polyvinylidene fluoride membrane (Bio-Rad, 1620177) and the indicated antibodies. The protein signals were detected using ECL western blotting detection reagents (PerkinElemer, NEL105001EA). The chemiluminescence bands were imaged under Amersham Imager 680 (GE Healthcare Life Sciences, USA). The bands were adjusted within the linear range and quantified using ImageJ2 (v2.3.0, NIH).

### Immunofluorescence microscopy

Cells grown on cover glasses were fixed with 4% paraformaldehyde (Thermo Fisher Scientific, 43368) in PBS for 15 min at room temperature. Subsequently, cells were permeabilized by 0.1% NP40 in PBS for 10 min at room temperature. After permeabilization, the cells were incubated with blocking buffer (2% BSA in cell staining buffer [4A Biotech, FXP005]) for 1 h at room temperature. Next, the cells were incubated with the indicated primary antibodies diluted in blocking buffer for 1 h at room temperature, and then incubated with fluorescent dye-conjugated secondary antibodies and DAPI in blocking buffer for 30 min at room temperature. Finally, the cells were mounted with Anti-Fade Fluorescence Mounting Medium (Abcam, ab104135). Images were acquired using a Zeiss LSM800 microscope (Zeiss, Germany) with a 63x 1.4 NA oil objective. The same acquisition parameters were used for a specific set of experiments. Z-series images were displayed as maximum Z-projections and orthogonal views processed with ImageJ2. The ratio of colocalization was determined by analyzing a total of 100 cells from three independent experiments. SIM images were obtained using the Zeiss Lattice SIM microscope (Zeiss, Germany) with a Plan-Apochromat oil objective lens 63x, 1.4 NA. Images were captured using PCO edge sCMOS camera (Acquisition pixel size, 63 nm at 63x objective) and G6 gride (23 μm). Data reconstruction was performed using the default 3D method in Zen Black software.

### GFP-Cleavage assay

U-2 OS cells were transiently co-transfected with *GFP-SEC61B* along with either scramble shRNA or *USP20* shRNA. After transfection, the cells were lysed using RIPA buffer (50 mM Tris-HCl, pH 7.5, 150 mM NaCl, 5 mM EDTA, 0.1% SDS, 1% Triton X-100 [Sangon Biotech, A110694-0100]), 0.5% sodium pyrophosphate, containing EDTA-free protease inhibitor cocktail [Bimake, B14002]) for 30 min at 4°C. The cell lysates were subjected to immunoblotting using antibodies against GFP, USP20, and ACTB.

### Microsome fraction assay

HEK293FT cells were washed three times with PBS buffer and then collected by centrifugation at 200 x g for 5 min. The cells were resuspended in three volumes of ice-cold sucrose buffer (10 mM HEPES, 250 mM sucrose, 2 mM MgCl_2_) containing EDTA-free protease inhibitor cocktail. The cells were mechanically lysed by passing through a 26-gauge needle, and the resulting cell lysate was centrifuged at 3,800 x g at 4 °C for 30 min. The supernatant was collected and subjected to a second centrifugation under the same conditions to remove any remaining debris. The resulting post-nuclear supernatant was centrifuged at 75,000 x g for 1 h at 4 °C using a TLA-55 rotor. The resulting microsome pellet was resuspended in microsome buffer (10 mM HEPES, 250 mM sucrose, 1 mM MgCl_2_, 0.5 mM DTT). Both the microsome and cytosolic fractions were subjected to immunoblotting using specific antibodies.

### Quantification and statistical analysis

All experiments were independently repeated at least three times. Image analysis and quantification were performed using ImageJ2. The image processing steps included background subtraction, smoothing, and conversion to binary masks through thresholding. The numbers in different groups were measured using the “analyze particle” function with fixed particle size and circularity parameters. Data were pooled from three independent experiments, with at least 100 cells counted. Error bars represent the standard error of the mean (SEM). Statistical analysis was performed using Prism 8 (GraphPad). The significance between two groups was determined using the Student’s *t*-test. The significance among multiple groups was determined using one-way ANOVA followed by Tukey’s multiple comparisons test. n.s., not significant; \**p* < 0.05, \*\**p* < 0.01, \*\*\**p* < 0.001, \*\*\*\**p* < 0.0001.

## Abbreviations

ACTB: actin beta
ADRB2: adrenoceptor beta 2
AMFR/gp78: autocrine motility factor receptor
ATG: autophagy-related
ATL3: atlastin GTPase 3
BafA1: bafilomycin A_1_
BECN1: beclin 1
CALCOCO1: calcium binding and coiled-coil domain 1
CCPG1: cell cycle progression 1
DAPI: 4’,6-diamidino-2-phenylindole
DUB: deubiquitinating enzyme
EBSS: Earle’s Balanced Salt Solution
FAM134B/RETREG1: (reticulophagy regulator 1)
FFAT: two phenylalanines [FF] in an acidic tract
GABARAP: GABA type A receptor-associated protein
GFP: green fluorescent protein
HMGCR: 3-hydroxy-3-methylglutaryl-coenzyme A reductase
IL1B: interleukin 1 beta
LIR: LC3-interacting region
MAP1LC3/LC3: microtubule associated protein 1 light chain 3
PIK3C3/Vps34: phosphatidylinositol 3-kinase catalytic subunit type 3
RB1CC1/FIP200: RB1-inducible coiled-coil 1
RFP: red fluorescent protein
RIPK1: receptor interacting serine/threonine kinase 1
RTN3L: reticulon 3 long isoform
SEC61B: SEC61 translocon subunit beta
SEC62: SEC62 homolog, preprotein translocation factor
SIM: super-resolution structured illumination microscopy
SNAI2: snail family transcriptional repressor 2
SQSTM1/p62: sequestosome 1
STING1/MITA: stimulator of interferon response cGAMP interactor 1
STX17: syntaxin 17
TEX264: testis expressed 264, ER-phagy receptor
TNF: tumor necrosis factor
UB: ubiquitin
ULK1/2: unc-51 like autophagy activating kinase 1/2
USP20: ubiquitin specific peptidase 20
USP33: ubiquitin specific peptidase 33
VAMP8: vesicle associated membrane protein 8
VAPs: vesicle-associated membrane proteins
VMP1: vacuole membrane protein 1
WIPI2: WD repeat domain, phosphoinositide interacting protein 2
ZFYVE1/DFCP1: zinc finger FYVE-type containing 1

## Acknowledgments

We thank Dr. Hong Zhang (Institute of Biophysics, Chinese Academy of Sciences), Professor Qiming Sun (Zhejiang University, China), and Professor Qing Zhong (Shanghai Jiao Tong University, China) for providing the various constructs. We thank Dr. Hong Zhang for providing valuable advice and critical reading of the manuscript. We thank the Image Core Facility and the Molecular and Cell Biology Core Facility in the School of Life Science and Technology at ShanghaiTech University for assisting with confocal microscopy and technical support.

## Author contributions

M.Z. and Y.L. conceived the project. M.Z. Z.W., Q. Z., Q. Y., C.Y, and Y.L. performed the experiments and analyzed the data. M.Z. and Y.L. wrote the manuscript.

