## Supplemental Materials for "USP20 deubiquitinates and stabilizes the ER-phagy receptor FAM134B to drive ER-phagy"

### Materials and Methods

#### *Plasmids*

*GFP-ULK1*, *GFP-BECN1* and *HA-ATG101* was described previously [1]. *MYC-ATL3* was described previously [2]. *GFP-WIPI2*, *GFP-SQSTM1* and *GFP-ZFYVE1* constructs were generated by inserting *WIPI2*, *SQSTM1* and *ZFYVE1* fragment into *pEGFP-C1* (Clontech, 6084-1). *pEGFP-ATG14/Atg14L* (21635; Tamotsu Yoshimori) was purchased from Addgene. *USP20-MYC-FLAG* was generated by inserting *USP20* fragment into *pCMV6-Entry* (OriGene Technologies, Inc., PS100001). *HA-ATG4B* was generated by inserting *HA-ATG4B* fragment into *pCDNA5* (Invitrogen, V6010-20). *pEnCMV-RTN3L(human)-3×FLAG-SV40-Neo* was purchased from MiaoLing Plasmid Platform (p30673). *FLAG-SEC62* and *FLAG-CCPG1* were generated by cloning the corresponding cDNA into the *pRK5-FLAG* vector. The construct for expression of *HA-TEX264* was kindly provided by Dr. Qiming Sun (Zhejiang University, China). The *PIK3C3-FLAG* was a kindly provided by Dr. Qing Zhong (Shanghai Jiao Tong University).

### Supplementary Figures

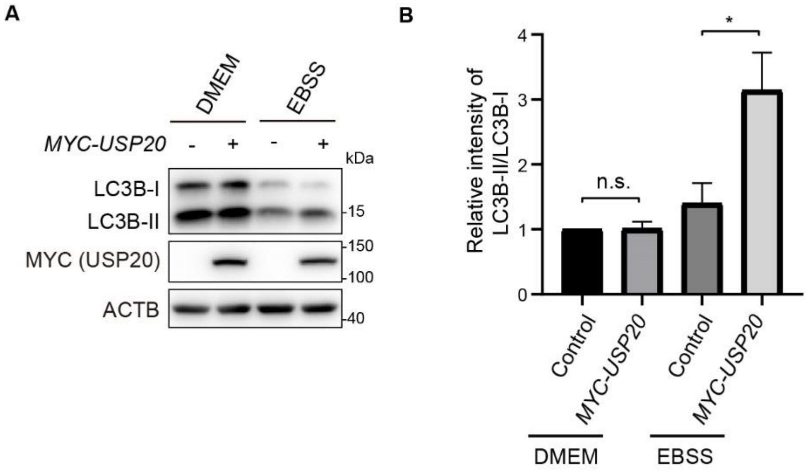


**Figure S1**. *USP20* overexpression results in increased ratio of LC3B-II to LC3B-I under starvation condition. (**A**) HeLa cells were transfected with the control vector or USP20. After 24 h, cells were treated with DMEM or EBSS for 4 h. The cell lysates were then subjected to immunoblotting with the indicated antibodies. ACTB was used as the loading control. (**B**) The relative intensity of LC3B-II/LC3B-I was quantified from (A). Statistical analyses were performed on data from three independent experiments. Error bars represent SEM. The significance levels are indicated as n.s., not significant, *$p < 0.05$ (one-way ANOVA with Tukey's test).

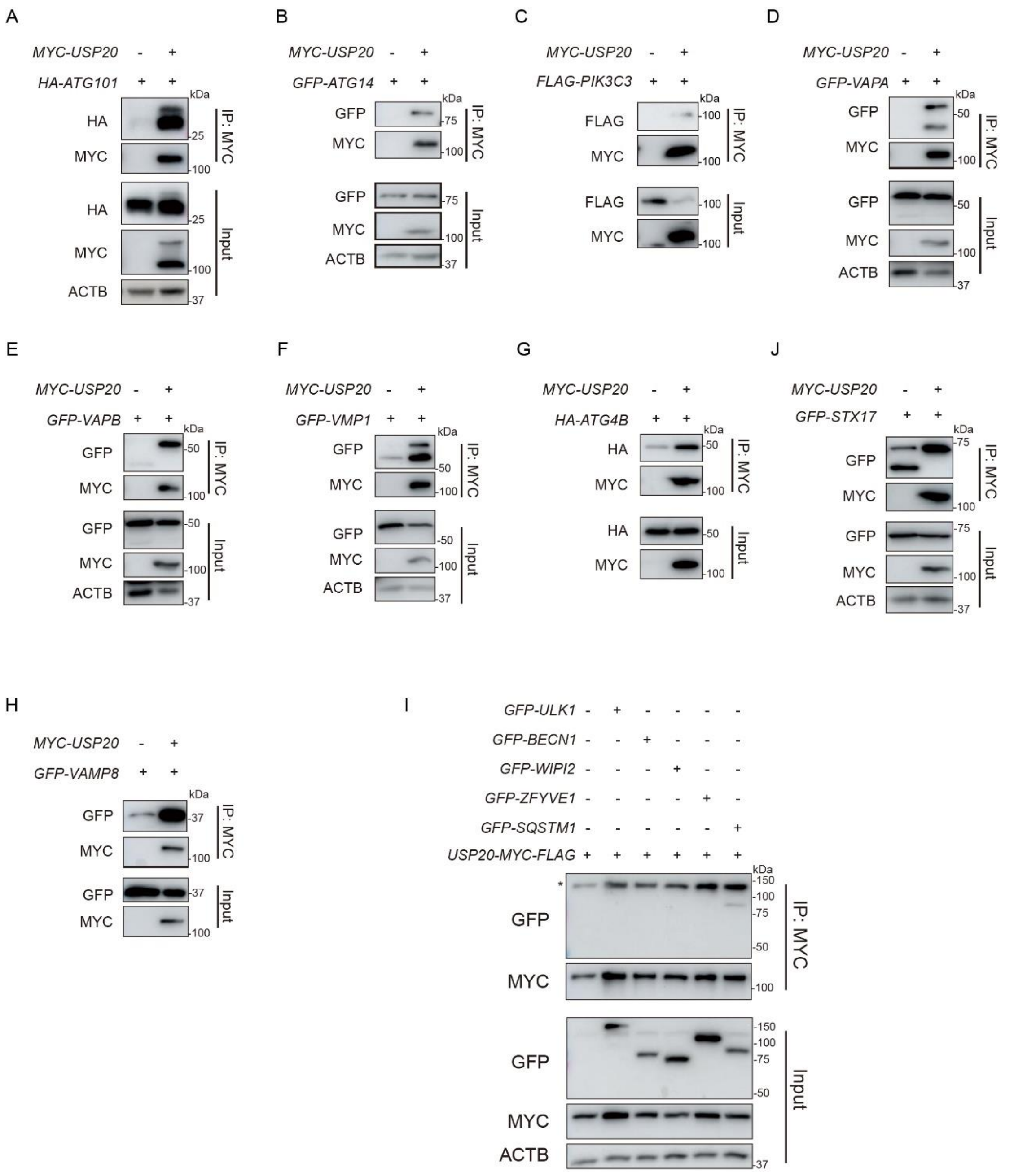


**Figure S2**. USP20 interacts with various autophagy-related proteins. (**A-I**) HEK293FT cells were transfected with the indicated constructs, along with either the empty vector or *MYC-USP20* (**A-H**). HEK293FT cells were transfected with *USP20-MYC-FLAG*, along with either the empty vector or GFP-tagged autophagy-related genes (**I**). MYC affinity isolation was performed using anti-MYC beads, and the isolated samples were subjected to immunoblotting analysis with the indicated antibodies. ACTB was used as the loading control.

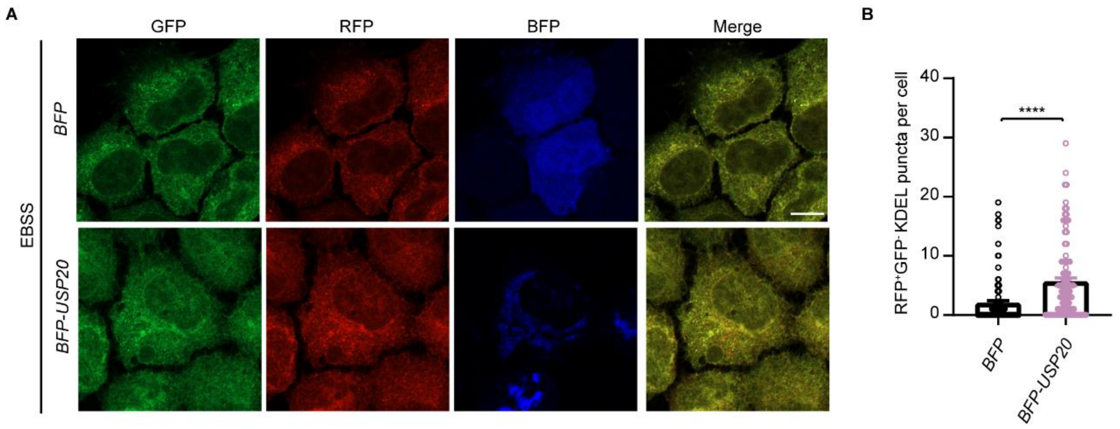


**Figure S3.** USP20 promotes ER-phagy. (**A**) HeLa cells stably expressing Tet-on *ssRFP-GFP-KDEL* were transfected with either the control BFP vector or *BFP-USP20*. After 24 h, doxycycline (DOX) was added to induce the expression of *ssRFP-GFP-KDEL*. After 24 h of induction, cells were treated with EBSS for 9 h. Cells were then fixed for fluorescence detection. Scale bar: 10 μm. (**B**) The number of RFP$^{+}$GFP$^{-}$ (red) KDEL puncta per cell was quantified from (A). Statistical analyses were performed on data from three independent experiments, with counts of more than 100 cells. Error bars represent SEM. The significance levels are indicated as ****$p$ < 0.0001 (one-way ANOVA with Tukey's test).

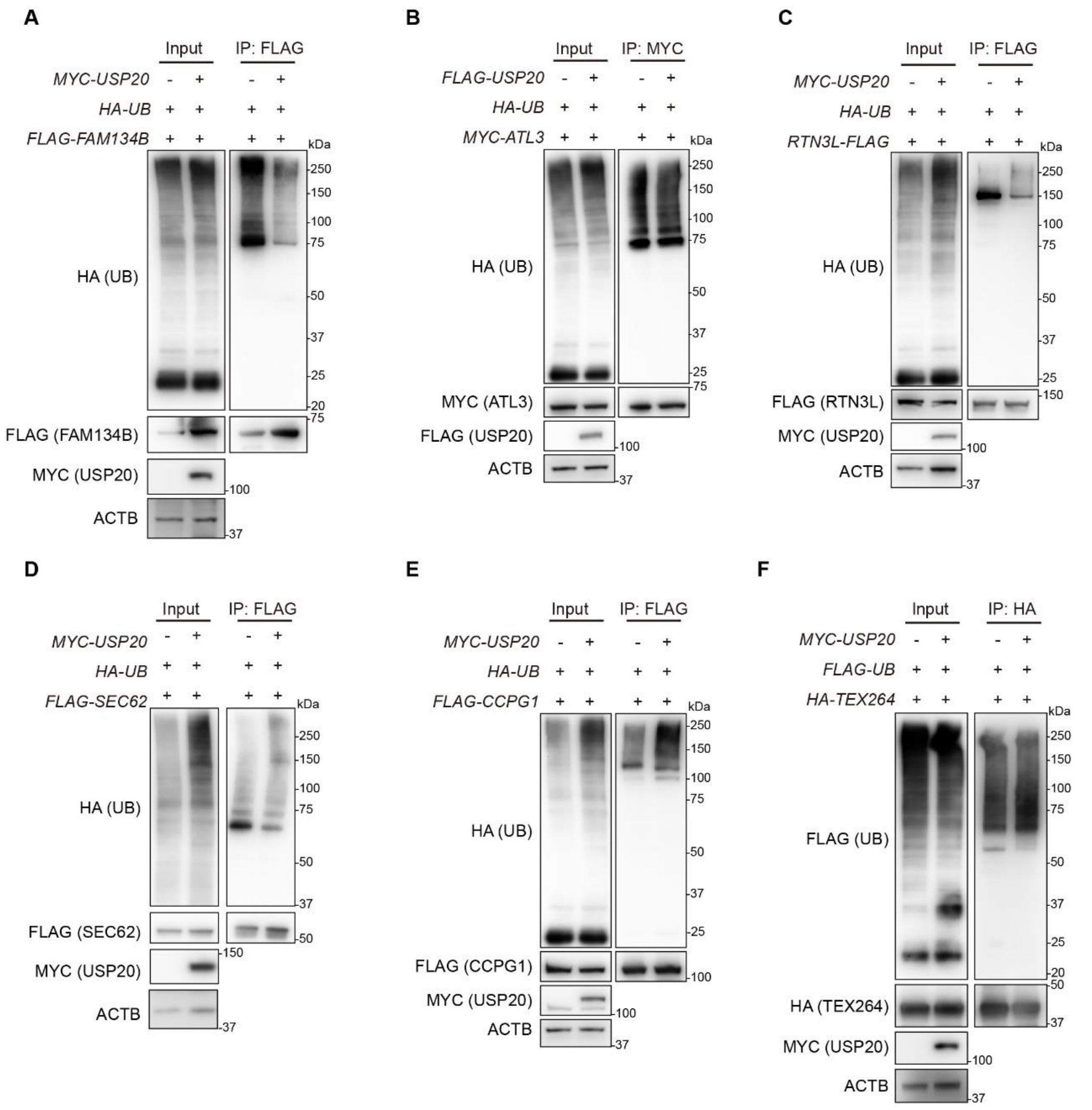


**Figure S4.** USP20 deubiquitinates FAM134B, but not other ER-phagy receptors. (**A-F**) HEK293FT cells were transfected with *FLAG-FAM134B* (**A**), or *MYC-ATL3* (**B**), or *RTN3L-FLAG* (**C**), or *FLAG-SEC62* (**D**), or *FLAG-CCPG1* (**E**), or HA-TEX264 (F), along with either the empty vector, *FLAG-USP20*, or *MYC-USP20*. Affinity isolation was performed using anti-MYC, FLAG, or HA beads under denaturing conditions, and the immunoprecipitated samples were subjected to immunoblotting analysis using specific antibodies to detect the levels of ubiquitination for each respective substrate. ACTB was used as the loading control in all the immunoblotting analyses.

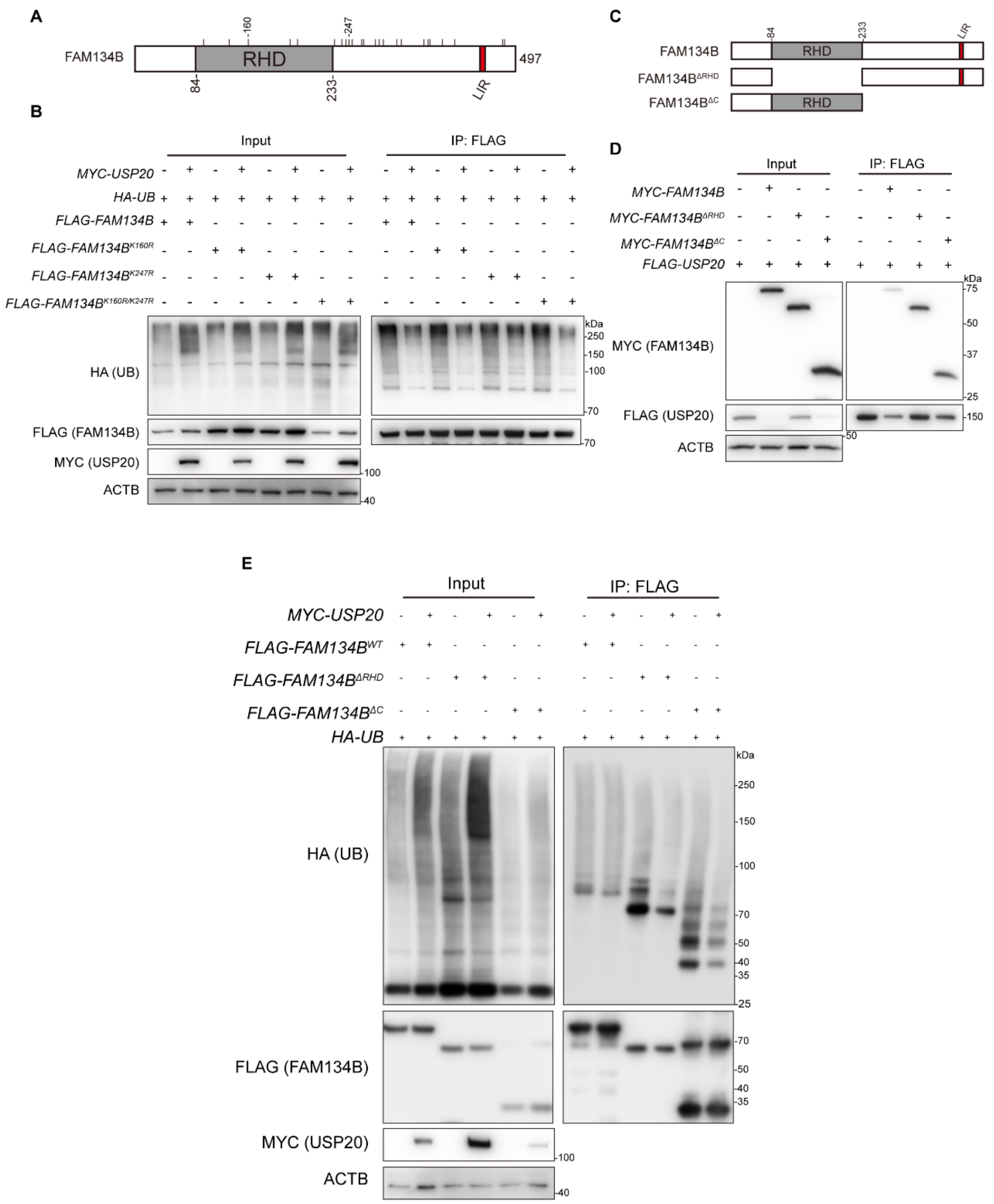


**Figure S5.** USP20 mediates the deubiquitination of FAM134B on multiple lysine residues. (**A**) The FAM134B protein sequence contains 24 lysine residues, including those within the Reticulon homology domain (RHD) and the C-terminal region. LIR: LC3-interacting region. (**B**) K160 and K247 of FAM134B are not the specific sites for USP20 deubiquitination. HEK293FT cells were transfected with *FLAG-FAM134B* wild-type, *FLAG-FAM134B*$^{K160R}$, *FLAG-FAM134B*$^{K247R}$, or *FLAG-FAM134B*$^{K160R/K247R}$, along with *HA-UB* and either the empty vector or *MYC-USP20*. FLAG affinity isolation was performed under denaturing conditions using anti-FLAG magnetic beads, and the immunoprecipitated samples were subjected to immunoblotting analysis using the specified antibodies to detect the levels of ubiquitination for FAM134B. (**C**) Schematic illustration of the FAM134B truncation mutants. (**D**) USP20 interacts with both the RHD and C-terminus regions of FAM134B. HEK293FT cells were transfected with *FLAG-FAM134B* wild-type, *FLAG-FAM134B*$^{\Delta RHD}$ or *FLAG-FAM134B*$^{\Delta C}$, along with USP20. FLAG affinity isolation was performed using anti-FLAG magnetic beads, and the immunoprecipitated samples were subjected to immunoblotting analysis using the specified antibodies. (**E**) USP20 mediates the deubiquitination of FAM134B on multiple lysine sites within different regions of the protein. HEK293FT cells were transfected with *FLAG-FAM134B* wild-type, *FLAG-FAM134B*$^{\Delta RHD}$ or *FLAG-FAM134B*$^{\Delta C}$, along with *HA-UB* and either the empty vector or *MYC-USP20*. FLAG affinity isolation was performed under denaturing conditions using anti-FLAG magnetic beads, and the immunoprecipitated samples were subjected to immunoblotting analysis using the specified antibodies to detect the levels of ubiquitination for FAM134B. ACTB was used as the loading control in all the immunoblotting analyses.

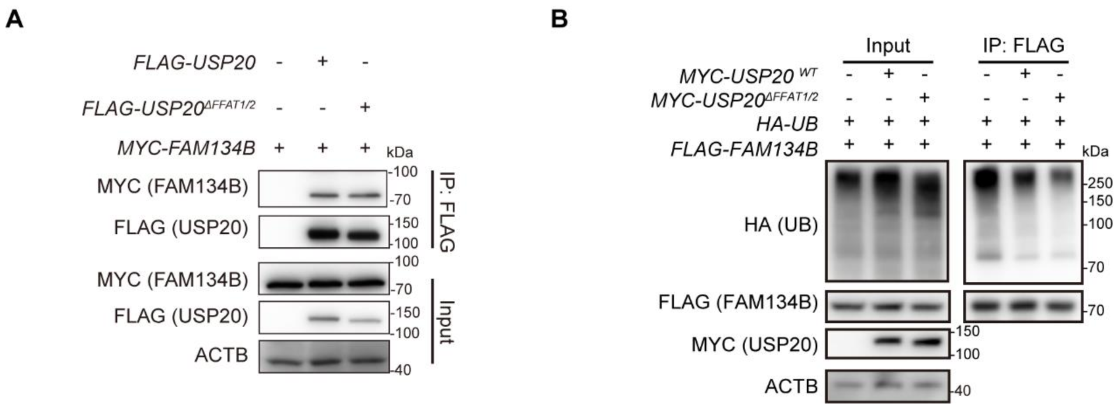


**Figure S6.** The interaction and deubiquitination of FAM134B by USP20 are not dependent on its FFAT motifs. (**A**) HEK293FT cells were co-transfected with either the empty vector or *FLAG-USP20* or *FLAG-USP20*$^{\Delta FFAT1/2}$, along with *MYC-FAM134B*. FLAG affinity isolation was performed using anti-FLAG magnetic beads, and the immunoprecipitated samples were subjected to immunoblotting analysis using the specified antibodies. ACTB was used as a loading control. (B) HEK293FT cells were co-transfected with either the empty vector or MYC-USP20 or MYC-USP20ΔFFAT1/2, along with FLAG-FAM134B and HA-UB. After 24 h of transfection, FLAG affinity isolation was performed under denaturing conditions using anti-FLAG magnetic beads, and the immunoprecipitated samples were subjected to immunoblotting analysis using the specified antibodies.

**Table S1.** List of oligonucleotides used in this study.

| **Target** | **Sequence (5’-3’)** |
|---|---|
| *USP20* KO sgRNA-1 | GTCAGTCGTGTGGGGTCACC |
| *USP20* KO sgRNA-2 | ATGTTGGCTGCGGAGAATCC |
| *VAPA* KO sgRNA-1 | TGAAGACTACAGCACCTCGC |
| *VAPA* KO sgRNA-2 | GGGTCAACTGTGACTGTTTC |
| *VAPB* KO sgRNA-1 | ACAGCGGAATCATCGATGCA |
| *VAPB* KO sgRNA-2 | ACTGACACTTCAGATATGGA |
| Scramble shRNA | CCTAAGGTTAAGTCGCCCTCG |
| *USP20* shRNA-1 | GCCCATCAGAAGATGAGTTCT |
| *USP20* shRNA-2 | CTATGTTGGCTGCGGAGAATC |
